# Joint Universal Modular Plasmids (JUMP): A flexible and comprehensive platform for synthetic biology

**DOI:** 10.1101/799585

**Authors:** Marcos Valenzuela-Ortega, Christopher French

## Abstract

Complex multi-gene plasmids can be built from basic DNA parts in a reliable and automation friendly way using modular cloning standards, based on Golden Gate cloning. However, each toolkit or standard is limited to one or a few different vectors, which has led to an overabundance of toolkits with varying degrees of compatibility. Here, we present the Joint Universal Modular Plasmids (JUMP), a vector design that overcomes the limitations of current toolkits by expanding the paradigm of modular cloning: all vectors can be modified using modular cloning in an orthogonal way using multiple cloning sites. This allows researchers to introduce any feature into any JUMP vector and simplifies the Design-Build-Test cycle of synthetic biology. JUMP vectors are compatible with PhytoBrick basic parts, BioBricks and the Registry of Standard Biological Parts, and the Standard European Vector Architecture (SEVA). Due to their flexible design, JUMP vectors have the potential to be a universal platform for synthetic biology regardless of host and application. A collection of JUMP backbones and microbial PhytoBrick basic parts are available for distribution.

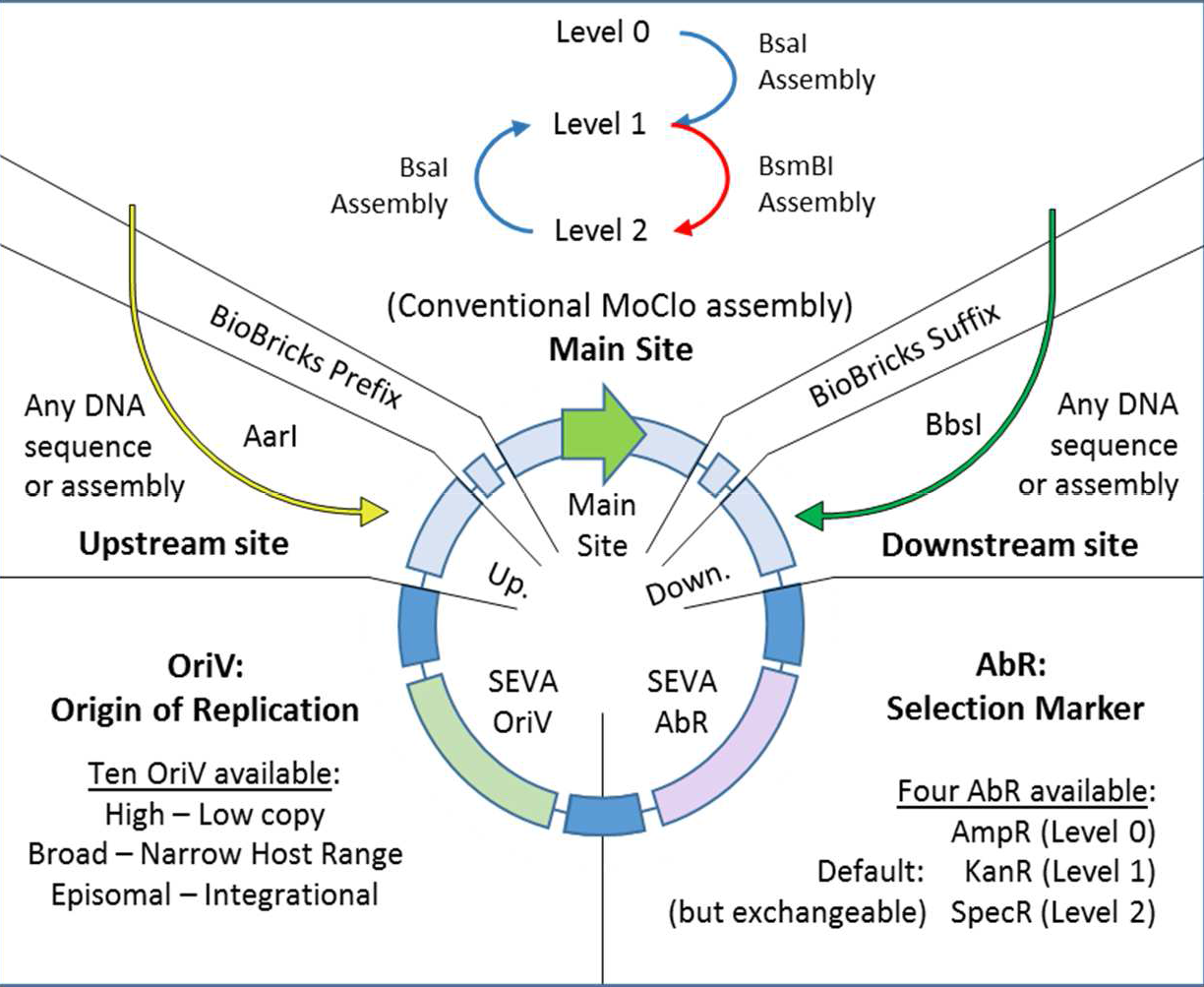

## Introduction

Synthetic biology aims to generate new-to-nature features in biological systems, approaching biology as an engineering field: simple DNA elements (or parts) are characterized and rebuilt to generate new genetic devices (1). Despite DNA synthesis becoming increasingly more affordable, the function of new DNA is hard to predict due to strong context dependency (2). Synthetic biology researchers have to re-iterate the design-build-test cycle several times until the new system is optimized or tuned and, consequently, DNA assembly standards that are robust, automatable and accept re-usable DNA parts are still necessary (3).

While Gibson assembly (4) and other overlap-based methodologies (as reviewed by Casini et al. (5)) are very efficient in assembling multiple DNA parts, standards based on Golden Gate cloning are the best fit for automation and part re-usability (5). In Golden Gate cloning (6), DNA parts are flanked by sites recognised by type IIS restriction enzymes, which cut outside the recognised site, leaving a user-designer fusion site. Parts are ligated in an ordered manner and restriction sites are removed during assembly, with assemblies of up to 24 parts (7). Different hierarchical standards based on Golden Gate have been published (8–17), collectively known as modular cloning (or MoClo) standards or toolkits. In MoClo standards, the product of one assembly level can be used as a part in the next level assembly, which is possible due to the use of compatible restriction sites and selection markers in the destination vector (Figure 1A). DNA parts need to be “domesticated”, a process that removes internal restriction sites and adds flanking restriction sites with the appropriate overhang, normally introducing the part into a plasmid for amplification and distribution. After a part has been domesticated, MoClo standards are PCR-independent, allow re-use of parts in different assemblies and allow building of complex constructs after multiple assembly rounds without PCR. This is a big advantage over overlap-based assembly systems, as PCR reactions can fail and are prone to introduce mutations and, consequently, impose the need for more extensive quality control of the constructs generated.

**Figure 1.**
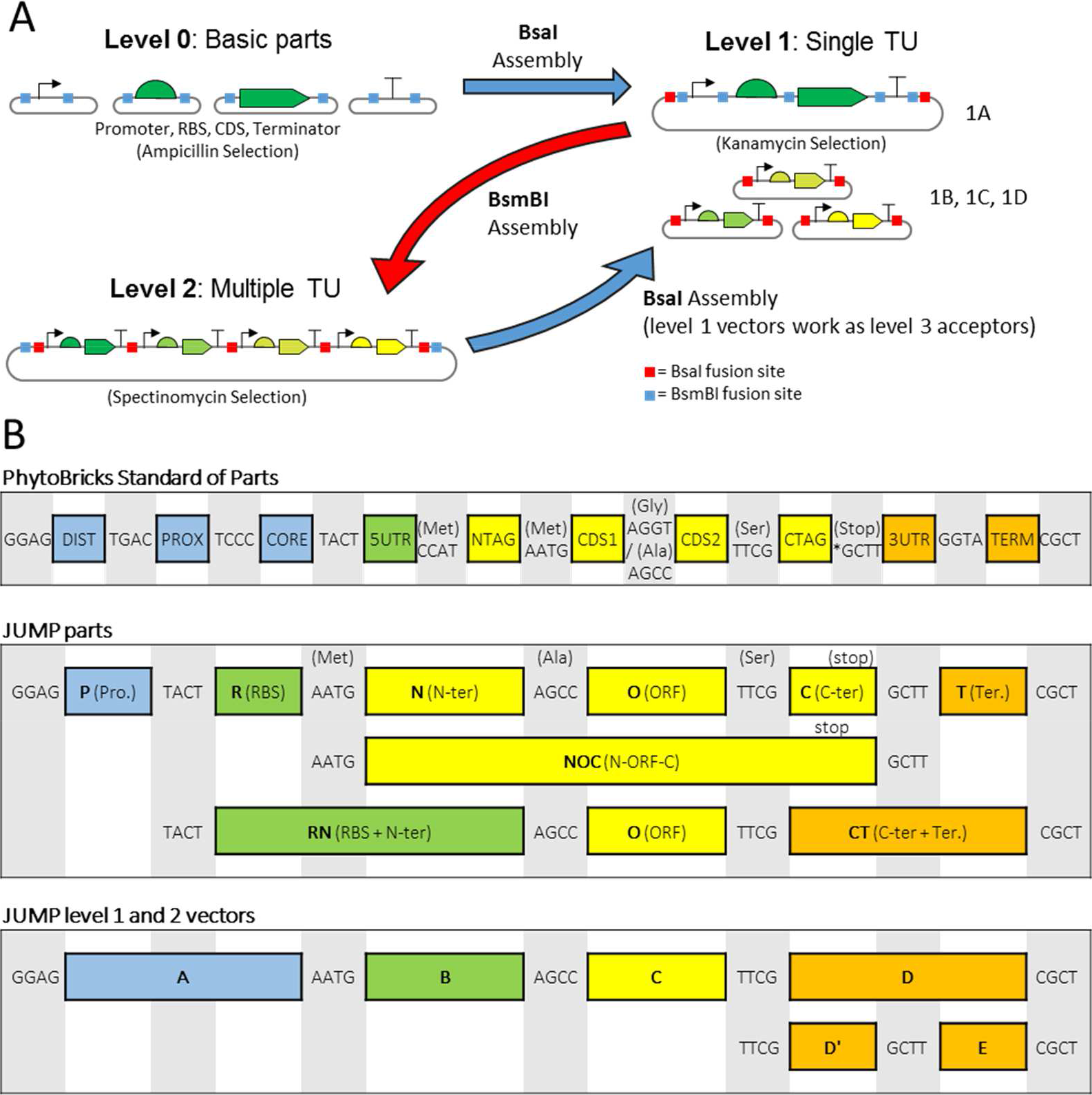
Modular cloning and PhytoBrick standardisation of parts. 1A) Basic parts are contained in level 0 vectors, which are assembled to form a single Transcription Unit in a level 1 vector. To simplify cloning and screening, the destination vector contains a different selection marker than insert-donor plasmids and a reporter gene replaced by the assembled TU. A level 2 vector can combine multiple level 1 assembly products, due to the use of an alternative type IIS restriction enzyme and selection marker between different assembly levels. In JUMP level 1 plasmids can be used as level 3 assembly destination vectors. 1B) PhytoBricks standard for fusion sites (18); format of JUMP parts provided in toolkit; fusion sites of level 1 and level 2 JUMP vectors.

The different MoClo standards differ in restriction enzymes used, fusion sites and the nature of the vectors, but the hierarchical design of modular cloning standards works similarly in the different standards (Figure 1A). Generally, each toolkit only offers one or a few vectors and for each host there will be an ideal vector containing specific genetic elements: origin of replication, selection marker and any other additional features needed. Therefore, vector toolkits are limited by the nature of their vectors. In addition to this vector constraint, the common MoClo pipeline imposes another limitation when a gene or sequence of interest (SOI) needs the presence of additional auxiliary DNA elements to be tested. The SOI is first assembled by itself and then assembled with the auxiliary elements. Multiple assemblies have to be done to study the SOI, even when the auxiliary elements are common for multiple SOI (e.g. transcription factors for inducible expression, dCas9 to test gRNAs, additional enzymes of a metabolic pathway when only one of them needs optimization…).

To allow compatibility of basic parts, multiple toolkits have adopted the PhytoBricks (18) common syntax that dictates that basic parts must be flanked by BsaI with specific fusion sites (Figure 1B). The PhytoBricks Standard was originally agreed among the plant synthetic biology community but is also used in standards compatible with bacteria (13,14,16).

Some MoClo toolkits include special features in their vectors for particular applications. For example, plant toolkits include LB/RB sequences flanking the cloning site to allow *Agrobacterium* mediated plant transformation (8,9,13,14,17) and some kits include unique flanking nucleotide sequences to allow combination of multiple TU via Gibson assembly (14). Another interesting additional feature is found in the EcoFlex toolkit (10). Some vectors have a “secondary module” to simplify pathway optimization. Multiple TU’s can be introduced in the secondary module (which uses an alternative type IIS nuclease) to then assemble 2 or 3 TU’s in the main site. Moore et al. showed that secondary sites increase assembly efficiency by decreasing the number of parts, which is highly desirable when generating libraries of clones. While the potential of secondary sites was demonstrated, it was limited by its design in EcoFlex plasmids, being restricted to special level 2 vectors that could only receive 2 or 3 TU in the main site, and the TU’s had to be pre-assembled into a level 2 vector before sub-cloning them into the secondary site.

Here we present “Joint Universal Modular Plasmids” (or JUMP), a vector standard designed to overcome the limitations of current modular cloning systems. JUMP vectors combine compatibility with PhytoBricks and BioBricks (19,20), with backbones based on the Standard European Vector Architecture (SEVA) (21,22). The SEVA repository is a large collection of origins of replication (OriV) and antibiotic selection markers (AbR) with a standardised format that allows simple exchange of vector elements (Figure 2A). Moreover, all JUMP vectors include two orthogonal secondary sites that can receive inserts from any MoClo level (Figure 2B). Thus, researchers can easily modify the vector chassis to reduce the steps needed for iterative assemblies and to give vectors new features (Figures 2C and 2D).

**Figure 2.**
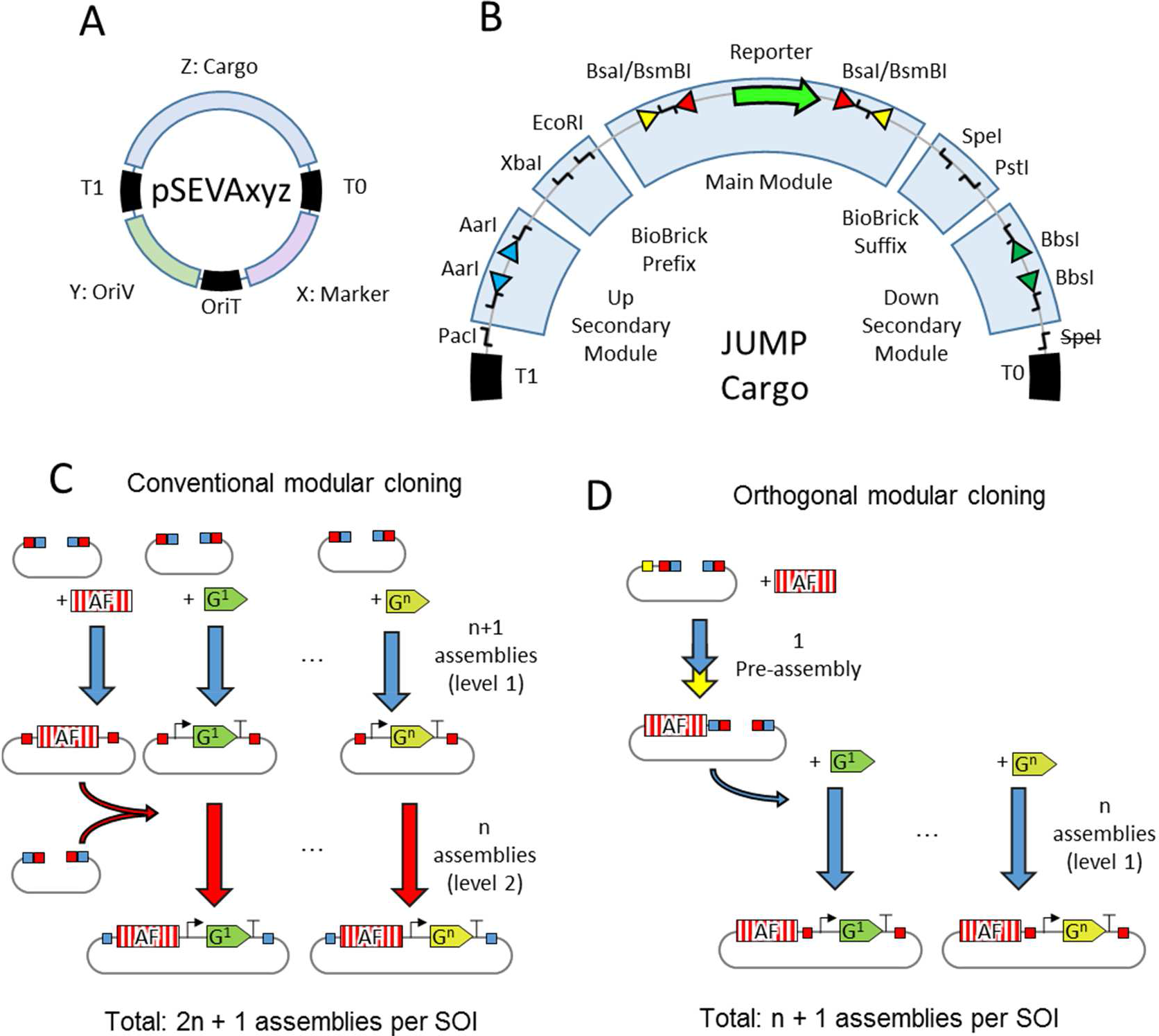
JUMP design and secondary sites. 2A) In SEVA (Standard European Vector Architecture) plasmids, three common short DNA sequences (black) flank three variable regions (coloured). Variable regions are the OriV (origin of replication), AbR (antibiotic selection marker) and “cargo” (any expression cassette). The invariable regions are two transcription terminators flanking the cargo (T1 and T0,) and origin of conjugation (oriT). Invariable regions also contain rare cutting sites, forbidden in the sequence of variable regions. 2B) JUMP is designed as special cargo of SEVA vectors to allow compatibility with future OriV’s and AbR’s of the collection. The cargo contains Upstream Module (outwards AarI); BioBricks prefix (XbaI, EcoRI); Main Module (a screening reporter gene flanked by outwards BsaI and inwards BsmBI for level 1, and vice-versa for level 2); BioBrick suffix (SpeI, PstI); and Downstream Module (outwards BbsI). SEVA’s canonical SpeI site was removed to allow BioBricks compatibility. 2C) Building constructs to test similar genes (G^1^ to G^n^) as sequences of interest (SOI) that depend on common auxiliary factors (AF) with conventional MoClo might require multiple assembly steps per SOI. D) Introduction of the auxiliary factors in vector chassis using orthogonal use of secondary modules.

## Methodology

### Strains and media

SEVA vectors were kindly provided by Victor de Lorenzo and his research group.

*Escherichia coli* JM109 (New England Biolabs) was used for transformations with assemblies. Selection and proliferation of strains was done with LB medium (10 g/L tryptone, 5 g/L yeast extract, 10 g/L NaCl) with the corresponding antibiotic depending on plasmid (100 μg/L carbenicillin for level 0 vectors, 50 μg/L kanamycin for level 1 vectors, spectinomycin at 50 μg/L for level 2 vectors. Vectors with chloramphenicol resistance were selected at 18 μg/mL. Homemade chemically competent cells were prepared based on the protocol of Chung et al. (23).

### Part domestication

We domesticated parts in the universal part-acceptor vector pJUMP18-Uac, which generates basic parts with any fusion site with overlapping BsaI and BtgZI. DNA fragments flanked by overlapping BsaI and BsmBI can be used for both part domestication into pJUMP18-Uac and directly for level 1 assembly (Supplementary Figure 2A). Promoters and terminators can be domesticated into the universal part acceptor or into the specialized acceptors pJUMP19-Pac (promoters) and pJUMP19-Tac (terminators), which allow part characterisation in *E. coli* as described in Results.

Parts must be free of BsaI and BsmBI to be compatible with JUMP level 1 and 2 assemblies, but we routinely removed AarI, BbsI, BtgZ sites to allow assemblies using those enzymes. We generated short parts with annealed oligonucleotides and longer parts either by PCR or by DNA synthesis (Integrated DNA Technologies). To domesticate parts via PCR and remove internal forbidden sites we followed a protocol based on the one described by Sarrion-Perdigones et al. (8), successfully domesticating parts with up to 5 internal sites (Supplementary Figure 2B). Parts are PCR-amplified between ends and internal forbidden sites as different sub-parts, which are assembled in the part-acceptor vector using BsmBI.

Most parts in the toolkit were built from BioBricks parts (19) or amplified from the genome of *E. coli* JM109. Terminators were adapted from the EcoFlex toolkit (10). The sequence of counter-selection CDS PheS A294G was obtained from Meyer et al. (24), the Lambda Red coding sequences from pKD46 and cI^ts^ (temperature-sensitive cI repressor) from pCP20 (25,26).

### Assembly conditions

Assembly conditions are explained in detail in Supplementary Methodology. Level 1 and 2 assemblies were performed similarly as in previous Modular Cloning publications: 20 femtomoles of all parts and destination vector were cyclically digested —with BsaI (New England Biolabs Ltd) at 37 °C or BsmBI (NEB) at 42 °C— and ligated with T4 DNA ligase (NEB) at 16 °C. Double-stranded oligonucleotide linker dummies were used to replace parts and reduce the number of inserts in some assemblies.

In the first step of the two-step assembly in secondary sties, equimolar amounts of the inserts are assembled without destination vector as in conventional modular cloning assemblies. Meanwhile, the destination vector is digested with AarI (Thermofisher) or BbsI-HF (NEB) and Shrimp Alkaline Phosphatase (NEB). In the second step, equimolar amounts of insert assembly and destination vector are ligated with freshly added T4 ligase enzyme and T4 ligase buffer (NEB) for one hour at 16 °C.

We screened assembly transformants similarly as in previously published modular cloning publications, including colony PCR and restriction digestion of plasmid DNA minipreps. We routinely checked for antibiotic resistance due to the serendipitous finding that insert-carrying plasmids co-transform at high frequency without the presence of their specific selecting antibiotic. We found that this also occurred using other MoClo toolkits and co-transformed colonies were sometimes more than 10% of the total (data not shown).

### Fluorescence measurements

Expression of sfGFP was measured to determine gene expression from different JUMP vectors. Absorbance and fluorescence were measured in a plate reader (FLUOstar Omega Microplate Reader, BMG Labtech). Overnight cultures were diluted 5 times and green fluorescence was measured in triplicate, with excitation and emission wavelengths of 485 and 520 nm, respectively, and was normalised by optical density measured at 600 nm. To characterise promoters and terminators, eGFP expression was measured in the same way as above, and relative fluorescence was normalised to that of the J23100 promoter.

The characterisation of constructs with mCherry as reporter was done by fluorescence with excitation and emission wavelengths of 584 and 610 nm.

## Results & Discussion

### Design and construction of JUMP Vector Backbones

Construction of JUMP vectors is explained in detail in Supplementary Figure 1. Forbidden restriction sites were removed from SEVA components as shown in Supplementary Tables 1 and 2. The JUMP ‘cargo’ (Figure 2B), consisting of upstream cloning module flanked by AarI sites, BioBrick prefix, main cloning module consisting of marker gene sfGFP flanked by BsaI and BsmBI sites, BioBrick suffix, and downstream cloning module flanked by BbsI sites, was then introduced. Level 1 (with Main Modules 1A, 1B, 1C, 1D) and level 2 vectors (with Main Modules 2A, 2B, 2C, 2D) were built as a “core set” using SEVA’s origin of replication #9 (pBRR322/ROP) with a medium copy number, and alternative vectors with different origins were built with the Main Module 1A and 2A as shown in Supplementary Table 3. This architecture allows researchers to build intermediate assemblies with the core set and then perform the final assembly with the oriV of choice. The default antibiotic markers are ampicillin/carbenicillin for level 0, kanamycin for level 1 and spectinomycin/streptomycin for level 2. Additional alternative vectors have been included in the toolkit with a different marker gene for screening (*lacZ’α*), and antibiotic marker (a broad host range chloramphenicol resistance gene from the shuttle vector pSEVA3b61 (27). Additionally, we included two pairs of vectors that can replace the vectors 1D (with 1D’ and 1E) and 2D (with 2D’ and 2E), which give the option of combining 5 inserts in level 2 and level 3 assemblies.

The nomenclature of JUMP vectors (Figure 3A) is conservative with SEVA’s to facilitate easy combination of new SEVA OriV and AbR with JUMP vectors. Of a vector “pJUMPxy-z”, x and y indicate the selection marker and origin of replication as indicated by SEVA rules, with the difference of forbidden sites being removed in the JUMP version (see figure 3B and 3C). The characteristics of the main modules —level and position— are indicated by the index z (Figure 3D). For example, vector pJUMP29-1A(sfGFP) has AbR #2 (Kanamycin), OriV #9 (pBBR322/ROP) and Main Module level 1 that would take position A in a level 2 assembly. Note that JUMP vectors do not strictly follow the SEVA rules and therefore are SEVA “siblings”, due to the SEVA native SpeI site being removed to make JUMP vectors compatible with BioBricks.

**Figure 3.**
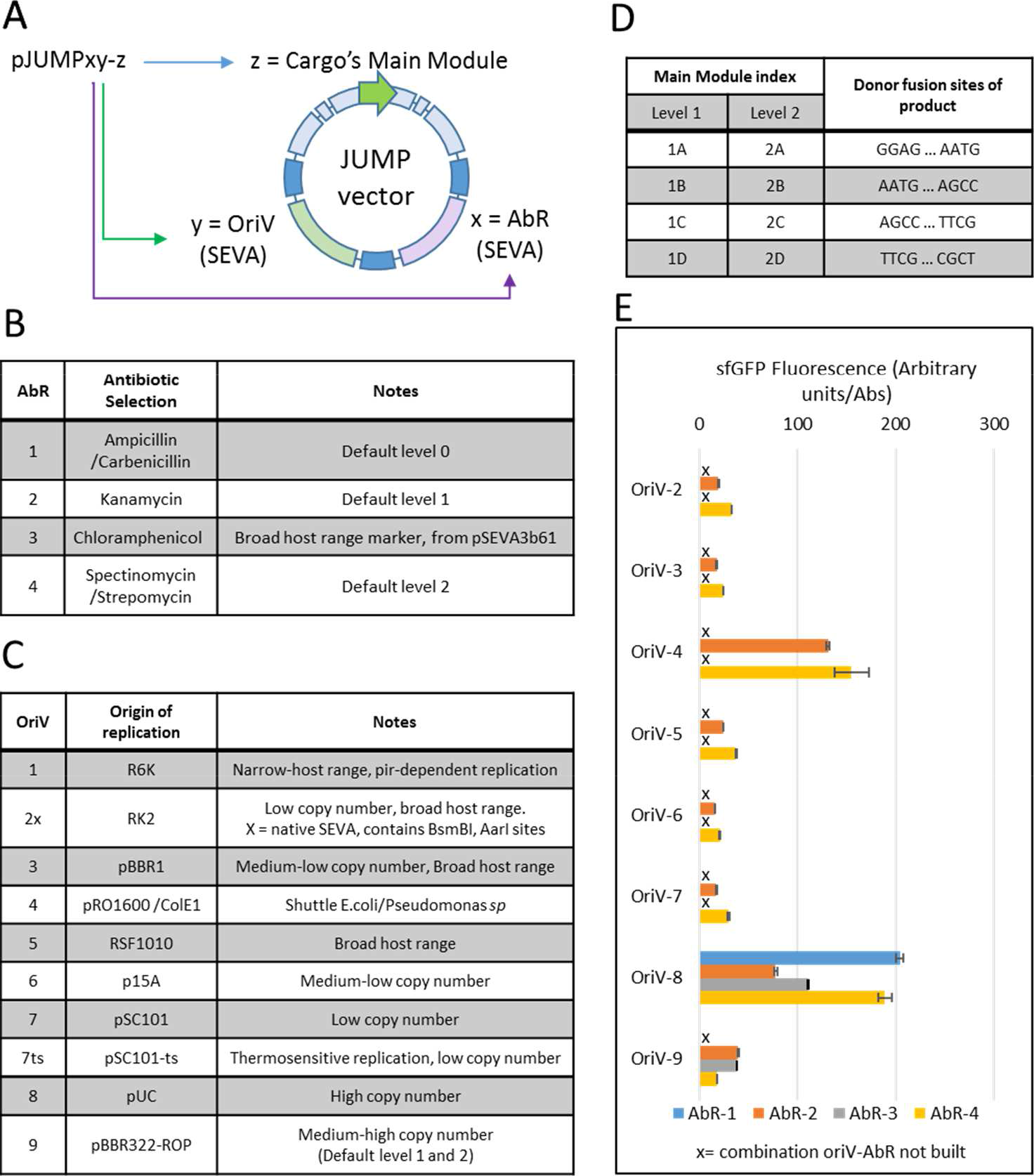
JUMP nomenclature and backbones. A) JUMP follows SEVA nomenclature for origins of replication (OriV) and selection marker (AbR), while cargo nomenclature is replaced by JUMP’s Main Module index. 4B) Antibiotic selection markers used in JUMP vectors. 4C) Origins of replication in JUMP vectors. 4D) Default main modules and their donor fusion sites. 4E) OD-normalised fluorescence of sfGFP cloning reporter in available combinations of OriV-AbR in the distributed JUMP toolkit. Error bars indicate standard deviation, n=3.

To allow compatibility with the widely used PhytoBrick basic parts, all acceptor fusion sites are constant (GGAG and CGCT) in all Main and Secondary Modules. In the Main Module, hierarchical conventional modular cloning works similarly as in current standards. PhytoBricks level 0 parts are assembled in any JUMP level 1 vector using BsaI (indicated by the removal of the marker gene); four level 1 products (1A, 1B, 1C, 1D) can be combined in any level 2 vector using BsmBI. As with Mobius (13) and Loop assembly (14,16), JUMP level 1 vectors can be used as assembly destinations for four level 2 products (2A, 2B, 2C, 2D) using BsaI. In addition to the compatibility with other toolkits, the use of constant acceptor sites offers the possibility to introduce sequences from any assembly level into either secondary site in any JUMP vector.

### Use of different JUMP vectors

Most MoClo toolkits only offer vectors with one origin of replication. Two recent toolkits offer multiple microbial OriV; the uLoop kit (16) offers four OriV with different copy number, and the MK toolkit (17) offers four bacterial broad-host range oriV as well as two for yeast. By contrast, the JUMP toolkit includes ten oriV, including broad-host range oriV, different copy-numbers and conditional vectors to allow chromosomal integration. Four replicative OriV allow its use with other hosts beyond *E. coli*, the *Pseudomonas* shuttle OriV #4 (pRO1600 /ColE1) and three broad host range vectors: origins #2x (RK2), #3 (pBBR1) and 5# (RSF1010). RK2 and pBBR1 are known to be replicative in a wide number of gram-negative bacteria, and the hyper-promiscuous RSF1010 has been additionally shown to replicate in some gram-positive bacteria and yeast species (28). We confirmed that JUMP vectors can be trans-conjugated from *E. coli* to *Pseudomonas putida* KT2440 (data not shown), using a tri-parental conjugation (29,30). We successfully conjugated *P. putida* with level 1 vectors with OriV #2, #3 and #5. Unexpectedly, the *Pseudomonas* shuttle OriV #4 did not yield any conjugants in our tests, although it contains no changes in the sequence from the original SEVA oriV which has previously been introduced into *P. putida* (31).

The availability of multiple vector backbones allows optimisation of the expression of the genetic construct. This is important as the characteristics of the vector strongly affect the expression of the gene it carries through context dependency (the effect of interactions of biological host complex environment and the function of the recombinant gene of interest) (10,11,15,32). Kim *et al* (32) showed the relevance of “tuning” the vector chassis by testing different constructs with multiple combinations of OriV and AbR’s. While the highest copy-number vectors were found to give a deleterious burden to host cells, other OriV’s showed different expression levels depending on the AbR they were paired with. The JUMP toolkit allows researchers to choose among ten different origins of replication and four antibiotic selection markers (Fig 3B and 3C). To show the effect of the vector chassis, we measured the expression of the constitutive sfGFP reporter in all empty vectors in the toolkit (Figure 3E). As expected, we found strong differences between different OriV and AbR. Expression was higher with the high-copy OriV #4 and #8, followed by the medium-copy #9, and the rest of OriV showed a lower expression. Surprisingly, expression from the medium-copy #6 was lower than expected. Jahn et al. (33) measured the copy number of different SEVA vectors and found oriV #6 to have a higher copy number than #7, although differences in expression and copy number could be due to a different strain being used.

The JUMP toolkit also offers conditional OriV’s to allow chromosomal integration. The SEVA OriV #1 (R6K) depends on the *pir* gene, not present in most *E. coli* strains. We inserted the vector pJUMP21-1A in *E. coli* JM109 (*pir*-) by introducing a 2kb homology sequence in a secondary site (obtaining colonies in the pir-*E.coli* strain JM109). Additionally, we have built a thermosensitive oriV (#7ts) by changing the Rep101 protein of SEVA’s oriV #7 (pSC101-based) with its thermosensitive counter-part from the plasmid pCP20 (25,26). After confirming that replication was temperature dependent (Supplementary Figure 5), we introduced a counter-selection marker (24) to allow curable chromosomal integration, which we used to generate a scarless transcriptional fusion of *gltA* in *E. coli* MG1655 to RBS and sfGFP parts from the toolkit (Supplementary Figure 6).

### Use of Upstream and Downstream Modules

While the Main Module enables modular cloning as with other MoClo systems, the advantage of JUMP resides in the presence of additional Golden Gate cloning sites in all vectors. The Upstream and Downstream modules use AarI and BbsI to introduce any sequence into the vectors, thus not disrupting the function of the Main module of any vector (level 0, 1 or 2). By being able to introduce any sequence in either secondary site of any vector, researchers can simplify the cloning steps needed to test the gene, genetic device or sequence of interest (SOI) (Figures 2C, 2D). Many recombinant SOI require other sequences to be present (auxiliary elements such as transcription factor for inducible expression, biosensor or reporter genes, Cas9/dCas9 for CRISPR, homology sequences for chromosomal integration, etc). JUMP allows such auxiliary elements to be introduced in a secondary site, thus reducing the number of assemblies when the auxiliary elements are common but multiple SOI will be assembled or require optimization.

The potential of Secondary Sites resides in how easily they can be used. By having the same receiving fusion sites, they are capable of receiving inserts assembled in the Main Module via a classic MoClo approach. A conventional Golden Gate assembly (one-pot and one-endonuclease) does not allow assembly of inserts into Secondary sites because the inserts are assembled with BsaI/BsmBI and this would cut the destination vector in the main site. JUMP basic parts were originally designed with overlapping BsaI and BtgZI sites, to allow double-enzyme assemblies as shown by Sarrion-Perdigones et al. (8); however, one-pot one-step assembly with BtgZI digesting level 0 inserts and AarI or BbsI digesting the destination vector showed very poor efficiency, possibly due to BtgZI remaining attached to the DNA and interfering with the assembly (34). Therefore, we use an efficient two-step assembly approach. (Figure 4A). In the first step the basic parts were assembled in the same conditions as for classic level 1 assemblies, and the destination vector was digested separately with AarI or BbsI for the upstream or downstream site, respectively, and dephosphorylated. In the second step, the digested backbone is ligated with the insert assembly mix. This was tested by assembling an mCherry TU from level 0 parts in the secondary sites of a level 1 destination vector. The transformation of the assembly resulted in more than 90% correct (mCherry+) colonies (Supplementary Figure 4). We expect that this approach can easily be automated, as there is no need to perform any purification steps or PCR reactions to prepare inserts, since inserts can come from the MoClo assembly pipeline without further modification. Additionally, AarI and BbsI sites do not have to be removed from inserts, as the restriction sites used for assembly on the main site (BsaI/BsmBI) are used on the sequence introduced in secondary modules. The two-step assembly can be extended to any insert and destination vector provided that the external fusion sites of the insert match the receiving fusion site of the destination vector (conserved for all JUMP vectors donor and acceptor sites), and the destination vector is selected with a different antibiotic than the insert donor vectors.

**Figure 4.**
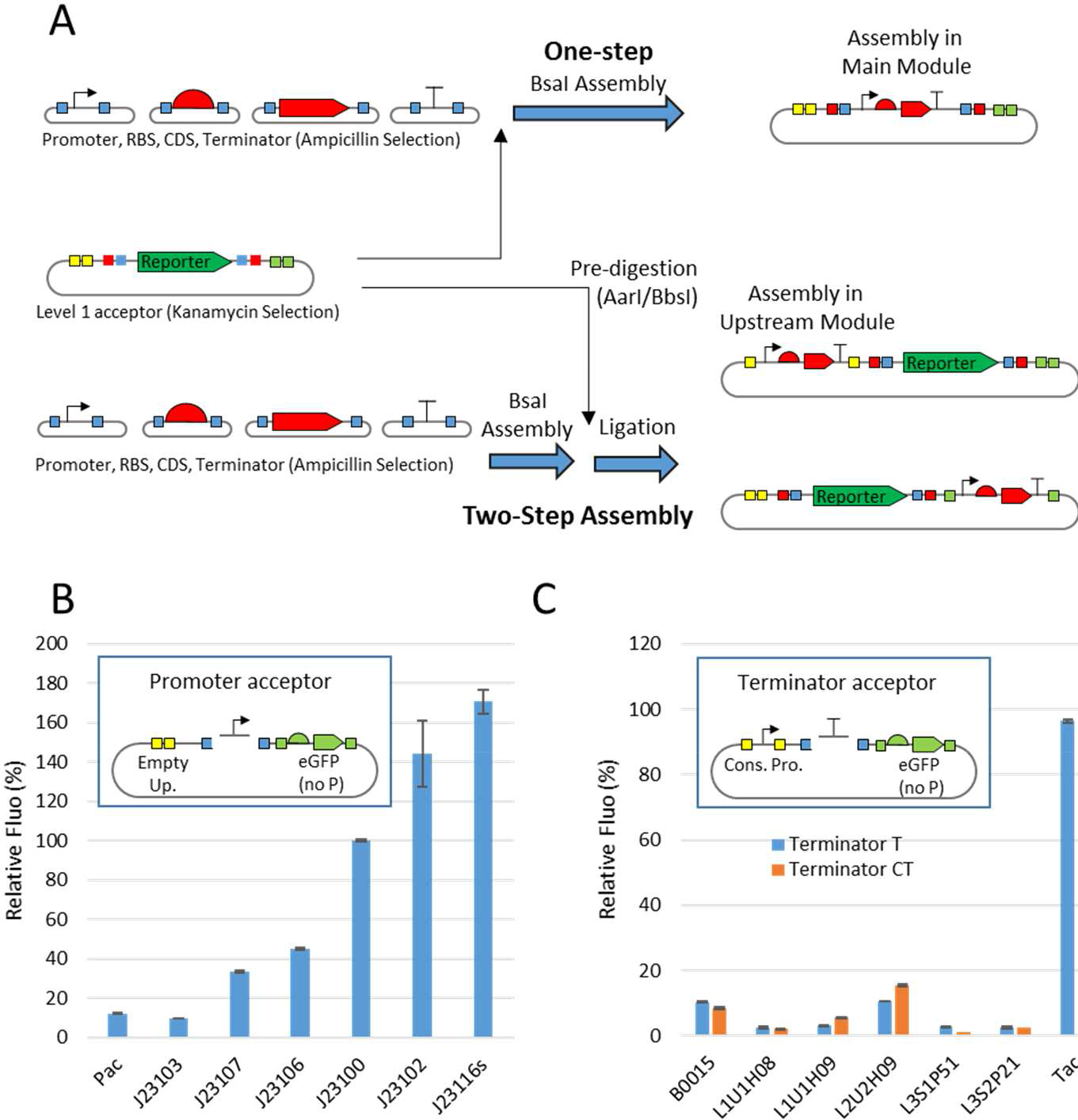
Use of secondary sites with two-step assembly. 4A) Two-step assembly works by assembling inserts first and then ligating the assembly reaction with the destination vector. 4B) Domestication and characterisation of promoters using level-0 promoter acceptor. 4C) Domestication and characterisation of terminators using level-0 promoter acceptor. T and CT terminators differ in 5’ end of the part (Figure 1B), with CT terminators including C-terminus stop codon. In 4B and 4C, Fluorescence was normalised to OD (600 nm) and is shown as % of that of the J23100 promoter. Error bars indicated standard deviation, n=3.

We used secondary sites to characterise promoters and terminators directly in the domestication (level 0) vector. We generated a level-0 promoter acceptor (pJUMP19-Pac) that reports the activity of the promoter by introducing eGFP without promoter in the Downstream site. During domestication of promoters, the eGFP indicated presence of the promoter and allowed characterization of the part (Figure 4B). We applied the same principle to domesticate and characterize terminators by introducing a constitutive promoter in the Upstream site, in such a way that the expression of the downstream eGFP would be disrupted by the presence of terminators in the main site (Figure 4C). Thus, part characterization can be done directly in the level 0 donor vector, rather than requiring level 1 or level 2 assemblies (10,11).

A second example shows how the screening of a combinatorial assembly can be simplified using the secondary site (Figure 5). We wanted to generate a library of the lambda phage repressor cI^ts^, which we domesticated from pCP20 (25,26). To characterise the clones of the library, a reporter with the promoter controlled by cI must be present. We reduced the number of assembly steps needed to one per cI^ts^ by building the library in a level 1 vector where we had previously introduced the reporter gene in the Upstream Module (Figure 5A). The clones from the library should vary in translation and stability of transcription factor, therefore we expected variability in the expression of the mCherry reporter at 30 °C and 37 °C, which we confirmed (Figure 5B).

**Figure 5.**
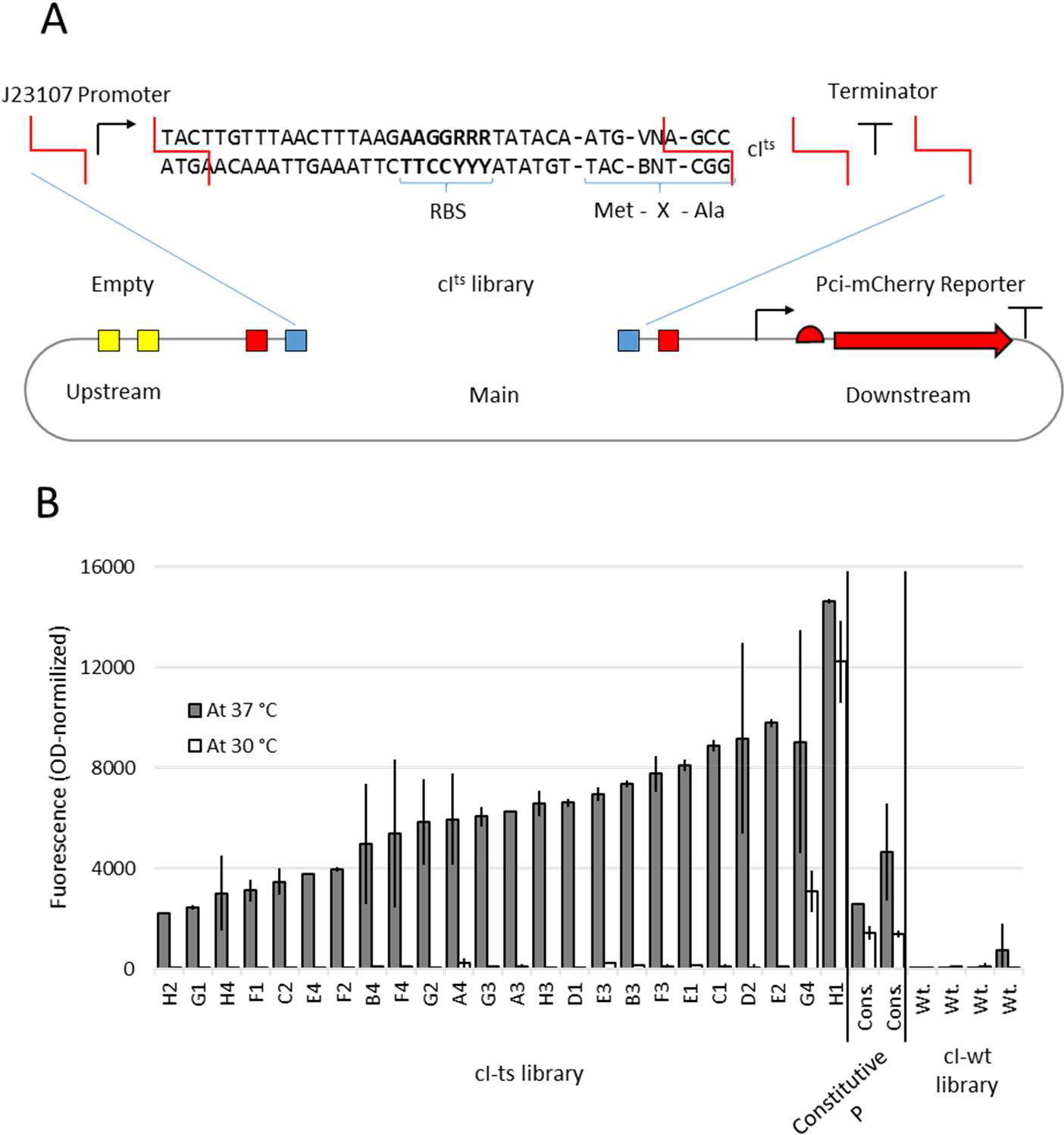
Use of secondary sites to screen variants of a transcription factor gene. 5A) The parts used for the construction of the cI^ts^ library were: a constitutive promoter, an RBS+N-terminus part, cI^ts^ CDS without N or C terminus, and C-terminus+Terminator. The RBS+N-terminus part was PCR-built from a degenerate oligonucleotide variable for the Shine-Dalgarno consensus and for the codon following the start ATG, thus giving the library diversity in translation initiation and half-life of protein (35). 5B) Colonies that were white at 30 °C were tested for expression of the mCherry reporter at 37 °C. As control, we analysed the fluorescence of two colonies from the library that were unrepressed at 30 °C (clones G4 and H1), two colonies with a constitutive mCherry gene, and four colonies from an equivalent library assembled with the wild-type (thermostable) cI CDS part.

It is important to note that virtually any sequence can be introduced in secondary sites. Some recent publications (13–15) have shown that new housekeeping elements normally present in the vector chassis (e.g. selection markers) can be introduced as assembly parts, thus increasing toolkit flexibility at the expenses of assembly complexity. By using secondary sites, any new feature can be introduced in JUMP vectors without increasing the complexity of later assemblies using the Main Module. Therefore, tailored plasmids can be built for any application: auxiliary elements to test the assembly in the main site, recombinase recognition sites, counter-selection markers, homology sequences for chromosomal integration, alternative origins of replication, etcetera.

### Toolkit Distribution

A collection of 96 vectors is available in Addgene (https://www.addgene.org/browse/article/28203402/). This collection includes a selection of PhytoBricks-compatible basic parts for synthetic biology applications in bacteria, a basic part Universal Acceptor, the level 1 and level 2 vector “core set”, and alternative vectors. We have removed type IIS restriction sites from all these basic parts and vectors, with the exception of the OriV #2x, (unmodified SEVA broad host range oriV #2/R2K, which is BsaI free but contains other forbidden sites).

### Conclusions

JUMP has been designed as a platform to allow MoClo assemblies to be introduced into any host and for any purpose. The capabilities of MoClo are combined with the vector flexibility of SEVA and multiple cloning sites. The toolkit generated allows any assembly level be done with ten different origins of replication. The vectors range from low to high copy number, as well as integrative vectors and they can be used with a very wide list of non-model microorganisms. Since they are based on SEVA, future OriV’s of the collection and AbR’s will be easily incorporated into JUMP vectors, and researchers can easily create new origins of replication and markers for the toolkit or use JUMP vectors with any PhytoBrick basic part. We have demonstrated that the Upstream and Downstream secondary sites expand the paradigm of modular cloning to allow orthogonal modification of the vector chassis.

## Supporting information

Sequences of JUMP toolkit

