## Supporting Information for "Joint Universal Modular Plasmids (JUMP): A flexible and comprehensive platform for synthetic biology"

#### Supplementary Information

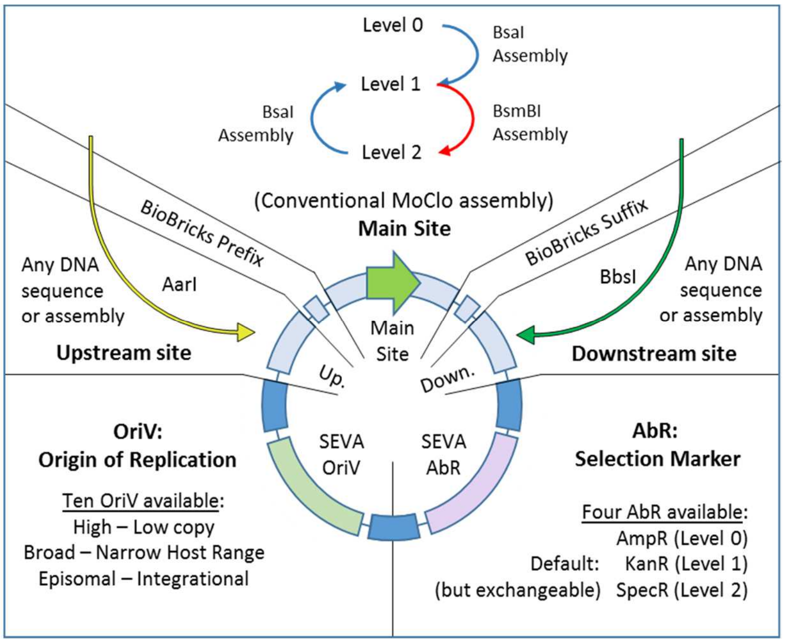


da

Authors: Marcos Valenzuela-Ortega[1]; Christopher French[1,2]

1. Centre for Systems and Synthetic Biology, School of Biological Sciences, University of Edinburgh, Roger Land Building, Alexander Crum Brown Road, Edinburgh EH9 3FF, UK

2. Zhejiang University-University of Edinburgh Joint Research Centre for Engineering Biology, International Campus, Zhejiang University, Haining, Zhejiang 314400,China

Supplementary Methodology.

Supplementary Table 1. Modification of SEVA selection markers

Supplementary Table 2. Modification of SEVA origins of replication

Supplementary Table 3. JUMP structural vectors in toolkit (Level 0, 1 and 2).

Supplementary Table 4. Basic parts in JUMP toolkit.

Supplementary Figure 1. Construction of JUMP toolkits.

Supplementary Figure 2. Domestication of parts

Supplementary Figure 3. Level 2 assembly.

Supplementary Figure 4. Two-step assembly results.

Supplementary Figure 5. Thermosensitivity of OriV #7ts and conditional integration.

Supplementary Figure 6. Chromosomal integration with conditional OriV #7ts

### Supplementary methodology

#### General materials and methods

Molecular biology reagents were obtained from New England Biolabs except for AarI (ThermoFisher). Oligonucleotides were obtained from Sigma-Aldrich.

For colony PCR screening GoTaq® G2 Green Master Mix from Promega Corporation was used.

Plasmid DNA used for assemblies was purified with the QIAprep Spin Miniprep Kit (QIagen) (plasmid), using 10 mL of LB overnight culture for medium and low copy number vectors, and 5 mL for high-copy number vectors (oriV #4, #8). DNA from PCR or digestions was purified with the QIAquick PCR Purification Kit (QIagen) or QIAquick Gel Extraction Kit (QIagen).

#### Oligo-linker preparation

Oligo-linker dummies were prepared by combining 5 µL of each oligonucleotide (100 µM), 34 µL of nuclease free water, 5 µL of T4 DNA ligase buffer and 1 µL T4 DNA polynucleotide kinase (PNK). They were incubated for 30 minutes at 37 ˚C, 95 ˚C for 5 minutes and cool down slowly to room temperature (0.2 ˚C per second if using a thermocycler) to allow annealing.

#### Conventional Modular Cloning (Main Module)

Parts are assembled in equimolar concentrations, with reaction and incubation conditions as explained below. BsaI is used for level 1 (and level 3 assemblies) and BsmBI for level 0 and level 2 assemblies.

Set-up (20 µL Reactions)

| Reagent | Initial Conc. | Vol. (µL) | Final Conc. |
|---|---|---|---|
| Backbone DNA | x | x | 20 fmol/20µL |
| Insert DNA | x | x | 20 fmol/20µL |
| T4 ligase buffer | 10 X | 2 | 1X |
| T4 DNA ligase (enzyme) | 400 unit/ µL | 0.25 | 100 u./20µL |
| BsmBI / BsaI-HF | 10/20 u./µL | 1 (or 0.5) | 10-20 u./20µL |
| $H_2O$ (nuclease free) | - | To give 20 µL | - |

Thermocycling incubation:

| "GG at 42" (BsmBI) | | |
|---|---|---|
| 42 °C | 15 min | |
| 42 °C | 3 min | x30 cycles |
| 16 °C | 3 min | |
| 55 °C | 15 min | |
| 80 °C | 5 min | |
| 10 °C | hold | |

| "GG at 37" (BsaI-HF) | | |
|---|---|---|
| 37 °C | 15 min | |
| 37 °C | 3 min | x30 cycles |
| 16 °C | 3 min | |
| 37 °C | 15 min | |
| 80 °C | 5 min | |
| 10 °C | hold | |

For difficult assemblies (higher number of parts, parts of bigger size) or when a higher colony number is desirable (combinatorial assemblies to generate libraries), the incubation protocol can be extended. Below there is a long protocol for overnight assemblies:

Long thermocycling incubation:

| "GG at 42 Super" (BsmBI) |
|---|

| "GG at 37 Super" (BsaI-HF) |
|---|

| | | |
|---|---|---|
| 42 °C | 15 min | |
| 42 °C | 5 min | X60 cycles |
| 16 °C | 5 min | |
| 55 °C | 60 min | |
| 80 °C | 5 min | |
| 10 °C | hold | |

| | | |
|---|---|---|
| 37 °C | 15 min | |
| 37 °C | 5 min | X60 cycles |
| 16 °C | 5 min | |
| 37 °C | 60 min | |
| 80 °C | 5 min | |
| 10 °C | hold | |

#### Two-step assembly (Secondary modules)

All JUMP vectors enable introduction of flanking sequences on either side of the Main Module. Conventional cloning allows introduction of any sequence using AarI (in the Upstream Module) or BbsI (in the Downstream Module). Moreover, because the receiver fusion sites of both secondary sites are the same as the ones used by the Main Module across all JUMP vectors (GGAG and CGCT), the sequences introduced in secondary sites can originate from other JUMP vector Main Modules. To maximise the utility of this design principle, we have developed a two-step assembly method to efficiently assemble TU from basic parts in either secondary site.

First step:

- Assemble inserts, without destination vector, using the methodology described in section 3.7, but increasing concentration to 40 femtomoles of each part per 20 µL reaction.
- We recommend using the long, overnight, protocol to increase efficiency.
- While the assembly reaction is running, pre-digest the destination vector:

| Upstream Pre-Digestion | Downstream Pre-Digestion |
|---|---|
| 200 femtomoles of destination vector | |
| 1 µL AarI | 1 µL BbsI-HF |
| 0.4 µL AarI's oligonucleotide 50X (see manufacturer's recommendations) | - |
| 2 µL AarI's Buffer 10X | 2 µL Cutsmart Buffer 10X |
| 0.5 µL Shrimp Alkaline Phosphatase | |
| Nuclease Free Water to a final volume of 20 µL | |

(Incubate at 37 °C for as long as the half-assembly is running and 20 minutes at 65 °C).

- A backbone pre-digestion can be used multiple times with different insert half-assemblies. Both first-step reactions can be stored frozen, but to store them for long periods, reactions should be purified with the PCR-purification kit.

Second step (mix and incubate).

- Ligate insert half-assembly and pre-digested destination vector:
    - 20 femtomoles of assembly product (10 µL of insert half-assembly reaction)
    - 20 femtomoles of pre-digested backbone (2 µL of reaction)
    - 2 µL of T4 ligase buffer 10X
    - 1 µL of T4 DNA ligase enzyme
    - Nuclease Free Water to a final volume of 20 µL
- Incubate for 1 hour at 16 °C

To select correct clones, transform *E. coli* (see note 1) with 1-2.5 µL of assembly reaction, and plate 10% of the assembly on LB agar containing the antibiotic corresponding to the destination vector.

Supplementary Table 1. Modification of SEVA selection markers

| AbR | Antibiotic Selection | Changes from original SEVA (bases from SEVA SwaI site) |
|---|---|---|
| 1 | Ampicillin /Carbenicillin | Unmodified. |
| 2 | Kanamycin | Unmodified |
| 3 | Chloramphenicol | PCR-amplified marker from pSEVA3b61 was introduced in SwaI/PshAI site of JUMP vectors. Modified from pSEVA3b61 to follow SEVA guidelines of format with primers: ccggttATTTAAATaacaaacgggattgacttttaaaaaagg + ttaattGACAAAAGTCccttcaaacttcccaaaggc). |
| 4 | Spectinomycin /Strepomycin | 1026C>T (BtgZI site) |

Supplementary Table 2. Modification of SEVA origins of replication

| OriV | Origin of replication | Changes from original SEVA (bases from SEVA FseI site) |
|---|---|---|
| 1 | R6K | Unmodified. |
| 2x | RK2 | Unmodified, contains 4 BtgZI, 2 AarI and 1 BsmBI sites. Deletion (56bp after position 2031) found in original SEVA OriV. |
| 3 | pBBR1 | 486C>T (BbsI site). |
| 4 | pRO1600 /ColE1 | Unmodified. |
| 5 | RSF1010 | 417G>T (BtgZI site), 1189G>T (BtgZI site), 1453G>A (BbsI site), 1962C>G (BbsI site). Deletion (2bp after position 3219) found in original SEVA OriV. |
| 6 | p15A | Unmodified. Deletion (1bp after position 599) found in original SEVA OriV. |
| 7 | pSC101 | Unmodified. |
| 7ts | pSC101-ts | 531C>A, 793C>A (from Rep101 present in pCP20) |
| 8 | pUC | Unmodified. Mutation (427T>A) found in original SEVA OriV. |
| 9 | pBBR322-ROP | 1029C>A (BsmBI site). |

Supplementary Table 3. List of JUMP structural vectors in Addgene toolkit (Level 0, 1 and 2).

| Well | PlasmidName | Vector level | Purpose |
|---|---|---|---|
| **A1** | pJUMP19-min2 | Level 0. | "Minimal cargo" vector. Allows introducing any part using BsmBI; but lacks cloning reporter. Can be used to generate new promoter/terminator acceptor vectors. |
| **A2** | pJUMP19-Pac | Level 0. | Promoter acceptor, Minimal cargo vector; with Downstream eGFP without promoter. Allows introducing any part using BsmBI. |
| **A3** | pJUMP19-Tac | Level 0. | Terminator acceptor Minimal cargo vector; with eGFP in downstream secondary site and constitutive promoter J23100 in upstream site. Allows introducing any part using BsmBI. |
| **A4** | pJUMP18-Uac | Level 0. | Universal Acceptor Plasmid. The fusion sites of the basic part generated are introduced during domestication. sfGFP reporter gene. High copy number oriV #8. |
| **A5** | pJUMP29-1A(sfGFP) | Level 1. Core set. | OriV 9 (pBBR322/ROP; medium copy number). |
| **A6** | pJUMP29-1B(sfGFP) | Level 1. Core set. | OriV 9 (pBBR322/ROP; medium copy number). |

| Well | PlasmidName | Vector level | Purpose |
|---|---|---|---|
| **A7** | pJUMP29-1C(sfGFP) | Level 1. Core set. | OriV 9 (pBBR322/ROP; medium copy number). |
| **A8** | pJUMP29-1D(sfGFP) | Level 1. Core set. | OriV 9 (pBBR322/ROP; medium copy number). |
| **A9** | pJUMP49-2A(sfGFP) | Level 2. Core set. | OriV 9 (pBBR322/ROP; medium copy number). |
| **A10** | pJUMP49-2B(sfGFP) | Level 2. Core set. | OriV 9 (pBBR322/ROP; medium copy number). |
| **A11** | pJUMP49-2C(sfGFP) | Level 2. Core set. | OriV 9 (pBBR322/ROP; medium copy number). |
| **A12** | pJUMP49-2D(sfGFP) | Level 2. Core set. | OriV 9 (pBBR322/ROP; medium copy number). |
| **B1** | pJUMP21-1A(sfGFP) | Level 1 alternative vector. | Alt. vector with Origin R6K (pir conditional). |
| **B2** | pJUMP22x-1A(sfGFP) | Level 1 alternative vector. | Alt. vector with Origin RK2 (broad-host-range). *CONTAINS one BsmBI and multiple BtgZI and AarI forbidden sites. |
| **B3** | pJUMP23-1A(sfGFP) | Level 1 alternative vector. | Alt. vector with Origin pBBR1 (medium-copy; broad-host-range) |
| **B4** | pJUMP24-1A(sfGFP) | Level 1 alternative vector. | Alt. vector with Origin pRO1600/ColE1 (E.coli - Pseudomonas shuttle) |
| **B5** | pJUMP25-1A(sfGFP) | Level 1 alternative vector. | Alt. vector with Origin RSF1010 (Broad-host-range) |
| **B6** | pJUMP26-1A(sfGFP) | Level 1 alternative vector. | Alt. vector with Origin p15A (medium copy number) |
| **B7** | pJUMP27-1A(sfGFP) | Level 1 alternative vector. | Alt. vector with Origin pSC101 (low copy number) |
| **B8** | pJUMP27ts-1A(sfGFP) | Level 1 alternative vector. | Alt. vector with Thermosensitive origin pSC101 (low copy number) |
| **B9** | pJUMP28-1A(sfGFP) | Level 1 alternative vector. | Alt. vector with Origin pUC (high copy number) |
| **B10** | pJUMP27ts-1A(lacZ) | Level 1 alternative vector. | With lacZ as alternative cloning reporter. Thermosensitive origin pSC101 (low copy number) |
| **B11** | pJUMP29-1A(lacZ) | Level 1 alternative vector. | With lacZ as alternative cloning reporter. OriV 9 (pBBR322/ROP; medium copy number). |
| **B12** | pJUMP29[dCas9]-1A(lacZ) | Level 1 alternative vector. | With lacZ as alternative cloning reporter.OriV 9 (pBBR322/ROP; medium copy number). Constitutive dCas9 in downstream secondary site. |
| **C1** | pJUMP27ts[mCherry]-1A(lacZ) | Level 1 alternative vector. | With lacZ as alternative cloning reporter. Thermosensitive origin pSC101 (low copy number). Constitutive mCherry in downstream secondary site. |
| **C2** | pJUMP29-1D'(sfGFP) | Level 1 alternative vector. | OriV 9 (pBBR322/ROP; medium copy number). |
| **C3** | pJUMP29-1E(sfGFP) | Level 1 alternative vector. | OriV 9 (pBBR322/ROP; medium copy number). |
| **C4** | pJUMP39-1A(sfGFP) | Level 1 alternative vector. | Level 1 vector with alternative selection marker (chloramphenicol). OriV 9 (pBBR322/ROP; medium copy number). |

| Well | PlasmidName | Vector level | Purpose |
|---|---|---|---|
| **C5** | pJUMP38-1A(sfGFP) | Level 1 alternative vector. | Level 1 vector with alternative selection marker (chloramphenicol). Origin pUC (high copy number). sfGFP cloning reporter. |
| **C6** | pJUMP41-2A(sfGFP) | Level 2 alternative vector. | Alt. vector with Origin R6K (pir conditional). |
| **C7** | pJUMP42x-2A(sfGFP) | Level 2 alternative vector. | Alt. vector with Origin RK2 (broad-host-range).*CONTAINS one BsmBI and multiple BtgZI and AarI forbidden sites. |
| **C8** | pJUMP43-2A(sfGFP) | Level 2 alternative vector. | Alt. vector with Origin pBBR1 (medium-copy; broad-host-range) |
| **C9** | pJUMP44-2A(sfGFP) | Level 2 alternative vector. | Alt. vector with Origin pRO1600/ColE1 (E.coli - Pseudomonas shuttle) |
| **C10** | pJUMP45-2A(sfGFP) | Level 2 alternative vector. | Alt. vector with Origin RSF1010 (Broad-host-range) |
| **C11** | pJUMP46-2A(sfGFP) | Level 2 alternative vector. | Alt. vector with Origin p15A (medium copy number) |
| **C12** | pJUMP47-2A(sfGFP) | Level 2 alternative vector. | Alt. vector with Origin pSC101 (low copy number) |
| **D1** | pJUMP47ts-2A(sfGFP) | Level 2 alternative vector. | Alt. vector with Thermosensitive origin pSC101 (low copy number) |
| **D2** | pJUMP48-2A(sfGFP) | Level 2 alternative vector. | Alt. vector with Origin pUC (high copy number) |
| **D3** | pJUMP49-2D'(sfGFP) | Level 2 alternative vector. | OriV 9 (pBBR322/ROP; medium copy number). |
| **D4** | pJUMP49-2E(sfGFP) | Level 2 alternative vector. | OriV 9 (pBBR+A15:D42322/ROP; medium copy number). |

Supplementary Table 4. JUMP Basic parts in Addgene toolkit.

| Well | Plasmid Name | Part type | Purpose |
|---|---|---|---|
| **D5** | pJUMP19-J23100_P | Basic Part P | Promoter (constitutive); J23100 |
| **D6** | pJUMP19-J23102_P | Basic Part P | Promoter (constitutive); J23102. |
| **D7** | pJUMP19-J23103_P | Basic Part P | Promoter (constitutive); J23103. |
| **D8** | pJUMP19-J23106_P | Basic Part P | Promoter (constitutive); J23106. |
| **D9** | pJUMP19-J23107_P | Basic Part P | Promoter (constitutive); J23107. |
| **D10** | pJUMP19-J23119s_P | Basic Part P | Promoter (constitutive); J23119 (1bp short on 3'). |
| **D11** | pJUMP19-T7pro_P | Basic Part P | Promoter (non-constitutive); T7 promoter; recognized by T7 polymerase |
| **D12** | pJUMP19-Pbad_P | Basic Part P | Promoter (non-constitutive); Pbad (I13453), arabinose induced; controlled by AraC. |
| **E1** | pJUMP19-Ptac_P | Basic Part P | Promoter (non-constitutive); Ptac; IPTG/lactose induced; controlled by LacI. |
| **E2** | pJUMP19-Ptet_P | Basic Part P | Promoter (non-constitutive); Ptet; induced by tetracycline/analogue; controlled by TetR |
| **E3** | pJUMP19-Pci_P | Basic Part P | Promoter (non-constitutive); Pci; controlled by cI. |
| **E4** | pJUMP19-Pars_P | Basic Part P | Promoter (non-constitutive); Pars; metal sensitive; controlled by arsR |
| **E5** | pJUMP19-Pmer_P | Basic Part P | Promoter (non-constitutive); Pmer; metal sensitive; controlled by merR |
| **E6** | pJUMP19-Pcue_P | Basic Part P | Promoter (non-constitutive); Pcue; metal sensitive; controlled by CueR |
| **E7** | pJUMP18-B0032_R | Basic Part R | RBS; B0032m Ribosome Binding Site. |

| Well | Plasmid Name | Part type | Purpose |
|---|---|---|---|
| **E8** | pJUMP18-B0033_R | Basic Part R | RBS; B0033m Ribosome Binding Site. |
| **E9** | pJUMP18-B0034_R | Basic Part R | RBS; B0034m Ribosome Binding Site. |
| **E10** | pJUMP18-RBS-pET_R | Basic Part R | RBS; pET vector Ribosome Binding Site. |
| **E11** | pJUMP18-RBS-pET-MV_RN | Basic Part RN | RBS+N-terminus; B0032m Ribosome Binding Site + Met-Val N-terminus start. |
| **E12** | pJUMP18-B0032-MV_RN | Basic Part RN | RBS+N-terminus; B0033m Ribosome Binding Site + Met-Val N-terminus start. |
| **F1** | pJUMP18-B0033-MV_RN | Basic Part RN | RBS+N-terminus; B0034m Ribosome Binding Site + Met-Val N-terminus start. |
| **F2** | pJUMP18-B0034-MV_RN | Basic Part RN | RBS+N-terminus; pET vector Ribosome Binding Site + Met-Val N-terminus start. |
| **F3** | pJUMP18-sfGFP_O | Basic Part O | ORF; sfGFP; fluorescent protein |
| **F4** | pJUMP18-amilCP_O | Basic Part O | ORF; amilCP; blue chromoprotein |
| **F5** | pJUMP18-araC_O | Basic Part O | ORF; araC; transcription factor |
| **F6** | pJUMP18-tetR_O | Basic Part O | ORF; TetR; transcription factor |
| **F7** | pJUMP18-arsR_O | Basic Part O | ORF; arsR; transcription factor |
| **F8** | pJUMP18-merR_O | Basic Part O | ORF; merR; transcription factor |
| **F9** | pJUMP18-lacI_O | Basic Part O | ORF; lacI; transcription factor |
| **F10** | pJUMP18-mCherry_O | Basic Part O | ORF; mCherry; fluorescent protein |
| **F11** | pJUMP18-cueR_O | Basic Part O | ORF; cueR; transcription factor |
| **F12** | pJUMP18-lacZ_O | Basic Part O | ORF; lacZ alpha-peptide; blue/white screening. |
| **G1** | pJUMP18-dCas9_O | Basic Part O | ORF; Catalytically dead mutant of the Cas9 from *Streptococcus pyogenes*. |
| **G2** | pJUMP18-cI_O | Basic Part O | ORF; cI; transcription factor from lambda phage. |
| **G3** | pJUMP18-cI-ts_O | Basic Part O | ORF; cI; transcription factor from lambda phage; thermosensitive. |
| **G4** | pJUMP18-pheS_NOC | Basic Part NOC | Complete ORF; PheS A294G; counter-selection marker; mutant phenylalanine tRNA synthetase. |
| **G5** | pJUMP18-lambdaRED_NOC | Basic Part NOC | Complete ORF; LambdaRED / λRED; Policistronic "NOC" part with coding regions for the three lambda re |
| **G6** | pJUMP18-Histag_N | Basic Part N | N-terminus; polyhistidine tag |
| **G7** | pJUMP18-Streptag_N | Basic Part N | N-terminus; Strep-tag II |
| **G8** | pJUMP18-Histag_C | Basic Part C | C-terminus; polyhistidine tag |
| **G9** | pJUMP18-Streptag_C | Basic Part C | C-terminus; Strep-tag II |
| **G10** | pJUMP19-L3S1P51_T | Basic Part T | Terminator; L3S1P51; synthetic terminator |
| **G11** | pJUMP19-B0015_T | Basic Part T | Terminator; B0015. Terminator. |
| **G12** | pJUMP19-L3S2P21_T | Basic Part T | Terminator; L3S2P21; synthetic terminator |
| **H1** | pJUMP19-L3S1P32_T | Basic Part T | Terminator; L3S1P32; synthetic terminator |
| **H2** | pJUMP19-L3S1P11_T | Basic Part T | Terminator; L3S1P11; synthetic terminator |
| **H3** | pJUMP19-L2U2H09_T | Basic Part T | Terminator; L2U2H09; synthetic terminator |
| **H4** | pJUMP19-L1U1H09_T | Basic Part T | Terminator; L1U1H09; synthetic terminator |
| **H5** | pJUMP19-L1U1H08_T | Basic Part T | Terminator; L1U1H08; synthetic terminator |
| **H6** | pJUMP19-B0015_CT | Basic Part CT | C-terminus + Terminator; B0015. C-termini STOP + Terminator. |
| **H7** | pJUMP19-L3S2P21_CT | Basic Part CT | C-terminus + Terminator; L3S2P21. C-termini STOP + terminator (synthetic) |

| Well | Plasmid Name | Part type | Purpose |
|---|---|---|---|
| **H8** | pJUMP19-L3S1P51_CT | Basic Part CT | C-terminus + Terminator; L3S1P51. C-termini STOP + terminator (synthetic) |
| **H9** | pJUMP19-L2U2H09_CT | Basic Part CT | C-terminus + Terminator; L2U2H09. C-termini STOP + terminator (synthetic) |
| **H10** | pJUMP19-L1U1H09_CT | Basic Part CT | C-terminus + Terminator; L1U1H09. C-termini STOP + terminator (synthetic) |
| **H11** | pJUMP19-L1U1H08_CT | Basic Part CT | C-terminus + Terminator; L1U1H08. C-termini STOP + terminator (synthetic) |
| **H12** | pJUMP18-cat_TU | Basic Part Whole TU | Codes for chloramphenicol resistance; including promoter and terminator. Broad host range. From Bacillus/E.coli shuttle pSEVA3b61. |

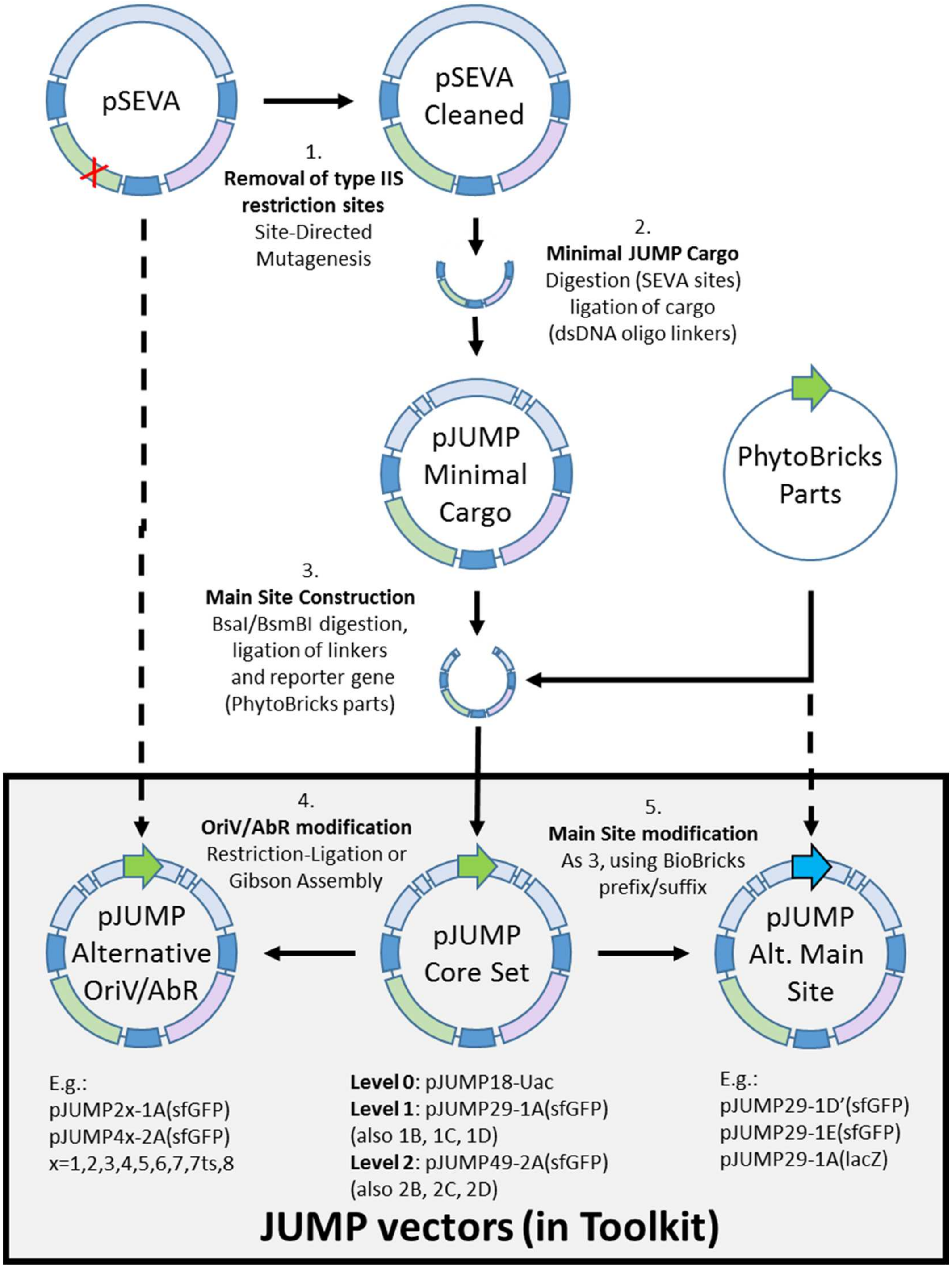


**Supplementary Figure 1. Construction of JUMP vectors.**

(1) For construction of the core set of vectors, forbidden sites were removed from SEVA vector using site-directed mutagenesis by blunt end ligation. (2) On the "cleaned" SEVA vectors, a minimal JUMP cargo was introduced between the SEVA PacI and SpeI sites. The minimal cargoes were built with short dsDNA linkers (oligonucleotides annealed and phosphorylated) that included Upstream and Downstream Module, Biobricks prefix and suffix, a minimal main module with only inwards BsaI (for building level 1 vectors) or BsmBI sites (for building level 0 and level 2 vectors). (3) We digested the minimal main module BsaI/BsmBI site to then ligate linkers (containing outwards BsmBI/BsaI and the fusion site specific for each vector) and a cloning reporter gene (built from digested Phytobricks parts). 4) Vectors with alternativer OriV or AbR were built using SEVA's

restriction sites or with Gibson assembly when SDM was used to remove forbidden sites. Thermosensitive OriV #7ts was built by combining the Rep101ts gene from pCP20 with SEVA's OriV #7 (pSC101 derivative).

The chloramphenicol marker AbR (#3) was cloned from pSEVA3b61 changing orientation to follow SEVA guidelines. (5) We used BioBricks sites to generate alternative main module (fusion sites or alternative reporter), using linkers and PhytoBricks parts or PCR fragments.

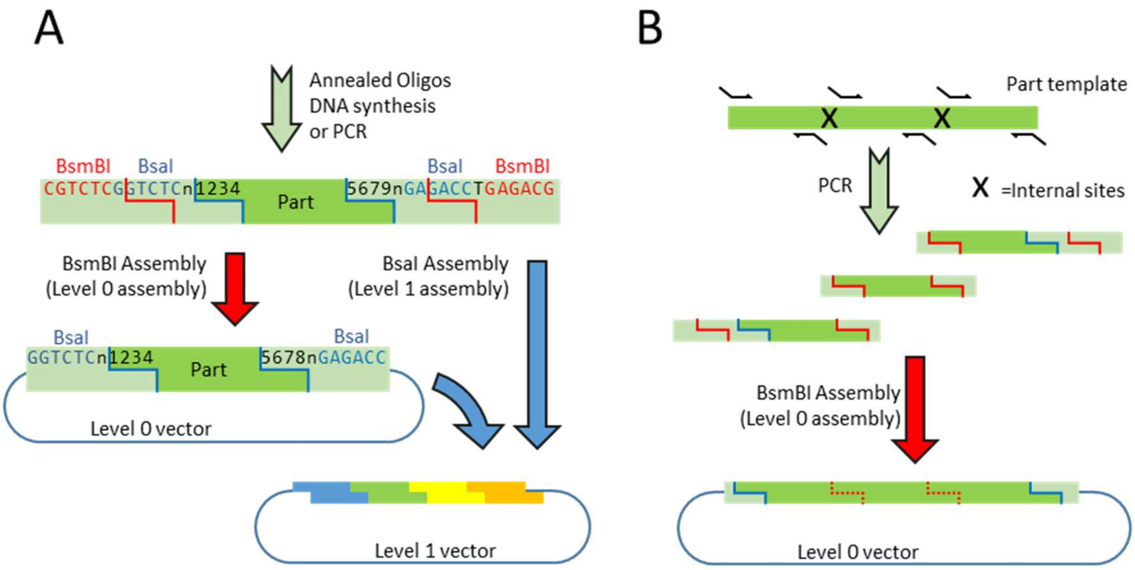


**Supplementary Figure 2. Domestication of parts.**

3A) Domestication of parts using a Universal Acceptor (such as pJUMP18-Uac). Linear fragments with overlapping BsaI and BsmBI can be used for both a level 1 assembly and for domestication Golden Gate Reaction to introduce the basic part into a level 0 vector. The part-specific fusion sites are represented as “1234” and “5678”. 3B) PCR-based domestication allows removal of internal forbidden sites during cloning. Parts are amplified as multiple sub-part fragments with primers that introduce silent single-point mutations removing internal sites and flanking BsmBI site in each sub-part fragment. BsmBI is used to assemble the sub-parts in the Universal Acceptor.

A

| Plasmid | Content |
|---|---|
| pNF49-2A (sfGFP) | Destination vector |
| pNF29-1A::J23106:His6:Chu2268 | B-glucosidase from *Cytophaga hutchinsonii*, constitutive |
| pNF29-1B::R00106:cenAo | Endoglucanase from *Cellulomonas fimi*, lactose promoter |
| pNF29-1C::J23106:Pdc | Pyruvate decarboxylase from *Zymomonas mobilis*, constitutive |
| pNF29-1D::J23106:Adh | Alcohol dehydrogenase from *Zymomonas mobilis*, constitutive |

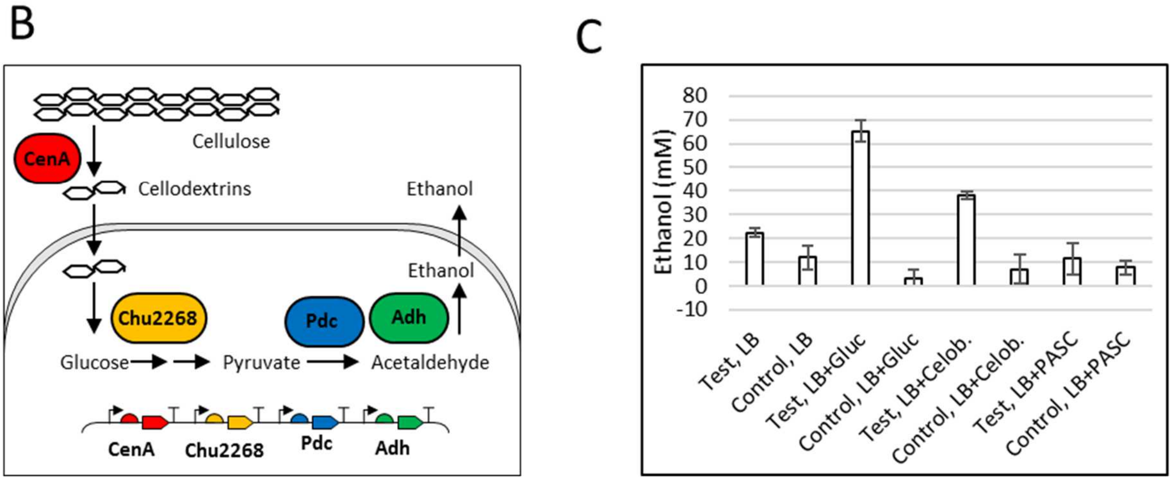


**Supplementary Figure 3. A level 2 assembly was performed to show the capability of JUMP vector for multi-gene vectors.**

A) Four level 1 assemblies were put together, each encoding an enzyme: two cellulases (CenA, Chu2268), a pyruvate decarboxylase (Pdc) and an aldehyde dehydrogenase (Adh). The basic parts coding for the enzymes were domesticated via PCR from plasmids containing these enzymes. The four genes were assembled correctly in pJUMP49-2A (confirmed by sequencing). B) The two cellulases allow *E.coli* to digest cellulose and to use it as carbon source (unpublished results). Adh and Pdc increase the ethanol yield in *E.coli* ([1]). C) Ethanol production of test strain (JM109+assembly) compared to control (JM109+empty pJUMp49-1A). Ethanol measured as explained by Lewicka et al ([1]): ethanol production was significantly increased from glucose and cellobiose. Even though we could not observe ethanol productionfrom cellulose, the activity of CenA was confirmed by digestion of carboxymethyl-cellulose (data not shown).

[1] Lewicka, A.J., Lyczakowski, J.J., Blackhurst, G., Pashkuleva, C., Rothschild-Mancinelli, K., Tautvaisas, D., Thornton, H., Villanueva, H., Xiao, W., Slikas, J. et al. (2014) Fusion of pyruvate decarboxylase and alcohol dehydrogenase increases ethanol production in Escherichia coli. ACS Synth Biol, 3, 976-978

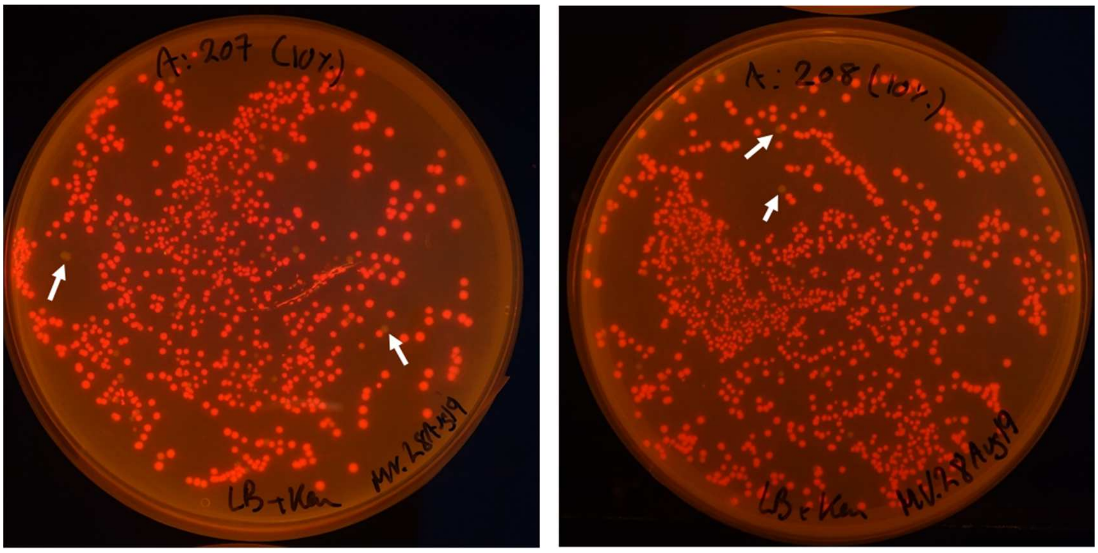

Upstream Downstream

**Supporting Figure 4. Two-step assembly of mCherry TU in secondary sites.**

Basic parts were combined to build a full transcription unit coding for a constitutive mCherry gene, which was assembled in the Upstream and Downstream Module of pJUMP29-1A(lacZ). Red fluorescence of colonies indicates correct assembly, while non-fluorescent colonies (as the ones pointed out by white arrows) are failed assemblies.

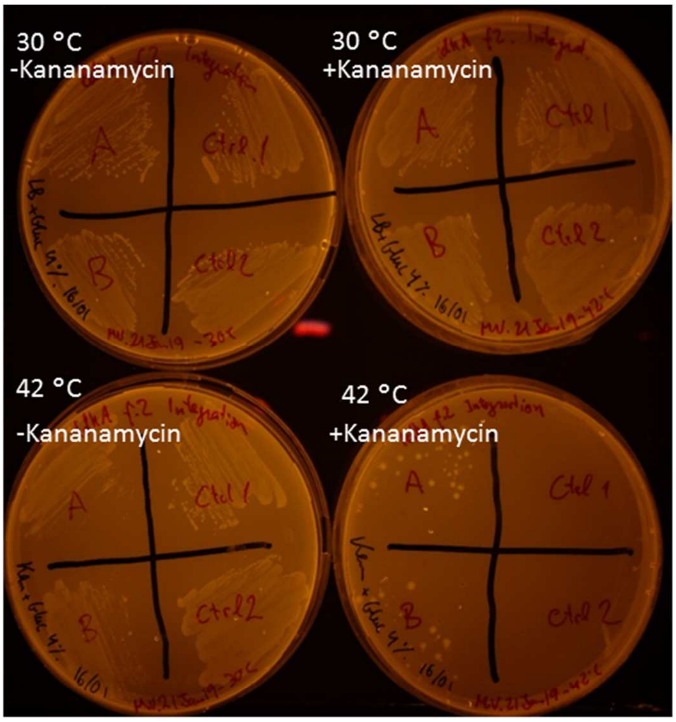


Ctrl 1 / 2:
pJUMP27ts-1A with no homology arms,

A and B:
pJUMp7ts-1A with homologous arms

| | Homology Arm | -Kanamycin | +Kanamycin |
|---|---|---|---|
| 30 °C | +HA | + | + |
| | No HA. | + | + |
| 42 °C | +HA | + | + |
| | No HA. | + | - |

**Supplementary Figure 5. Thermosensitivity of OriV #7ts and conditional integration.**

Conditional integration with OriV #7ts. Plasmid pJUMP27ts-1A(lacZ) (with thermosensitive replication) was tested in *Escherichia coli* MG1655 for plasmid stability and integration. While *E. coli* with the empty plasmid was able to grow carrying the plasmid at 30 °C or in the absence of kanamycin, only when homology sequence of 2 kb was inserted in the plasmid it was only to show colonies in the presence of kanamycin at 42 °C.

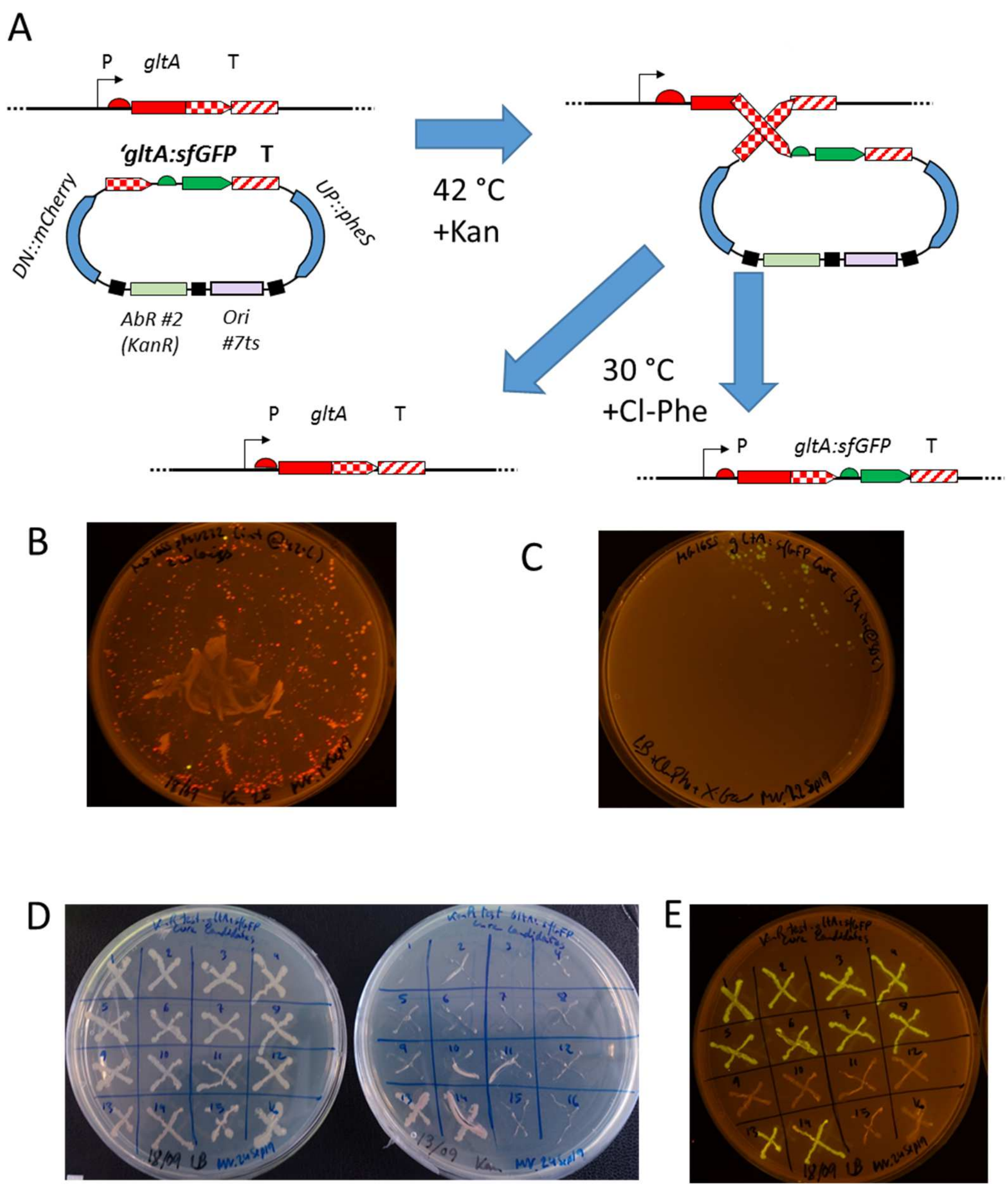


**Supplementary Figure 6. Chromosomal integration with conditional OriV #7ts.**

A) We introduced an mCherry reporter and pheS A294G counter-selection marker (1) in in upstream and downstream modules of pJUMP27-1A(lacZ), respectively. In the Main Module, we assembled a 400 bp left homology arm amplified from the end of the *E.coli gltA* gene, RBS and sfGFP parts, and 400 bp right homology arm amplified from the 400 bp following *gltA. E.coli* MG1655 was transformed and vector integration was forced by maintaining kanamycin selection at 42 °C. We incubated a GFP+ colony for 3 hours to promote a second recombination and selected curation by streaking cells in the presence of 8mM 4-chloro-L-phenylalanine. B) Plate with integrants. Homology of *pheS* with chromosomal wild-type gene led to high rate of off-site integrants (mCherry+ GFP-). C) Cure of a GFP+ integrant lead to circa 50% GFP+ and 50% GFP- colonies from recombination in right and left homology arms, respectively. D) Cured colonies were tested for kanamycin sensitivity. Left plate: LB only, right plate: LB+kanamycin. Samples: 1-8: GFP+ colonies, 9-12: GFP- colonies, 13-14: integrant pre-cure (KanR) control, 15-16: Wild-type (KanS) control. E) GFP expression of cured colonies (same samples). Correct integration was confirmed by PCR using flanking primers (data not shown).
